# scAGG: Sample-level embedding and classification of Alzheimer’s disease from single-nucleus data

**DOI:** 10.1101/2025.01.28.635240

**Authors:** T. Verlaan, G. A. Bouland, A. Mahfouz, M.J.T. Reinders

**Affiliations:** Delft University of Technology

## Abstract

Identifying key cell types and genes in Alzheimer’s Disease (AD) is crucial for understanding its pathogenesis and discovering therapeutic targets. Single-cell RNA sequencing technology (scRNAseq) has provided unprecedented opportunities to study the molecular mechanisms that underlie AD at the cellular level. In this study, we address the problem of sample-level classification of AD using scRNAseq data, where we predict the disease status of entire samples from the gene expression profiles of their cells, which are not necessarily all affected by the disease. We introduce scAGG (single-cell AGGregation), a sample-level classification model that uses a sample-level pooling mechanism to aggregate single-cell embeddings, and show that it can accurately classify AD individuals and healthy controls. We then investigate the latent space learnt by the model and find that the model learns an ordering of the cells corresponding to disease severity. Genes associated with this ordering are enriched in AD-linked pathways, including cytokine signalling, apoptosis, and metal ion response. We also evaluate two attention-based models that perform on par with scAGG, but entropy analysis of their attention scores reveals limited interpretability value. As scRNAseq is increasingly applied to large cohorts and cell-level disease association annotations do not exist, our approach provides a way to classify phenotypes from single-cell measurements. The yielded cell- and sample-level severity scores may enable identification of AD-associated cell subtypes, paving the way for targeted drug development and personalized treatment strategies in AD.

Code is available at: http://github.com/timoverlaan/scAGG

## Introduction

Recent advances in RNA sequencing technology allow measurement of gene expression at the individual cell level, through single-cell RNA sequencing (scRNAseq), which has revealed the heterogeneous landscape of cell types and allowed researchers to build cell atlases of various healthy tissues.^1,2^ It has also been used to study the cell-type-specific effects of diseases. In the context of Alzheimer’s disease (AD), one of the first single-cell studies identified disease-associated microglia (DAM) that have the potential to restrict neurodegeneration.^3^ However, analysis of such scRNAseq data introduces a new fundamental problem, because in bulk RNAseq, where the RNA expression of an entire piece of tissue is measured, the metadata of a sample contains information about the disease status of the individual and can be directly related to the measured gene expression of the sample. This approach has, for example, allowed the identification of three distinct AD subtypes that exhibit different combinations of dysregulated pathways.^4^ But in scRNAseq, the disease metadata is still at the sample-level, whereas the measured gene expression is now at the cell-level. We refer to this as the *sample-level classification problem in single-cell data*.

Several methods have been proposed to predict the disease status of single-cell data,^5–8^ but they focus on the task of cell-level classification. That is, they label each cell individually, according to the disease status of the sample they belong to. Alzheimer’s disease, however, is a highly heterogeneous disease that progresses over an extremely long time course, and it has been shown that different types of cells are affected differently and to different extents.^9–11^ So, if we label the individual cells according to the disease state of the sample they belong to, this may actually be incorrect at the cell-level.

We argue that the task of sample-level classification should be considered, where the sample is classified as a whole, based on the gene expression of all cells. This way, we do not explicitly enforce the wrong label upon unaffected cells in a diseased sample. The main challenge here is the variability in number of cells between samples, and the permutation invariant nature of the input data. That is, a solution should be able to handle samples of different sizes, and the order of the cells should not matter. To our knowledge, only a few methods directly consider the task of sample-level classification. However, to do so, they either rely on heavy down-sampling to get a fixed number of cells per sample,^12,13^ or use an unsupervised approach to learn sample-level embeddings that are not specifically optimised for classification.^14,15^

In this work, we propose scAGG (single-cell AGGregation), a machine learning model that performs the task of sample-level classification, that does not require a fixed number of cells per sample and directly uses the cell-level data through application of a pooling mechanism. This three-part architecture embeds individual cells, aggregates them into a sample-level embedding through a sample pooling mechanism, and classifies the sample according to disease class, all trained in an end-to-end fashion. We investigate several parametrizations of this architecture, including attention-based modules which allow for interpretability of the model, and we compare the performance of these models to several baseline approaches. scAGG is computationally efficient and scales linearly in the number of genes and cells.

## Materials and Methods

### Data processing

#### ROSMAP

10X chromium single-nucleus gene expression data was collected from 427 dorsolateral prefrontal cortex (DLPFC) samples of brain donors of the ROSMAP^16^ project. For information on the processing of reads and the generation of count matrices, we refer to the original study.^17^ Further processing of the counts was performed similar to the standard SCANPY^18^ approach for single-cell data: we discard genes detected in fewer than 200 cells and remove cells with less than 200 genes detected or more than 5% mitochondrial gene expression. Next, we normalise each cell to add up to 10, 000 transcripts (CP10k) and log-transform (log1p). We also remove a distinct cluster of 10 outlier samples, which were dominated by a specific cell type (RELN and CHD7 expressing excitatory neurons). Finally, we select only the 1, 000 most highly variable genes (HVGs), using the standard procedure in Seurat.^19^ The resulting dataset contains 339 samples, with a total of 2, 037, 478 cells.

From the complete set of individuals, we select the 78 most extreme cases of AD and the 61 least cognitively impaired controls, leaving 200 intermediate donors. This was done following the approach proposed in,^20^ where the Braak stage and CERAD scores, a neuropathological assessment and cognitive test respectively, are used as additional constraints to select only the most severe AD cases and healthiest controls as follows:

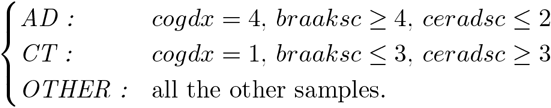

#### SeaAD

For the SeaAD^21^ replication dataset, we used the same processed steps as for ROSMAP. As only 959 genes overlapped with the top 1000 HVGs in ROSMAP, these genes were selected when training the model on ROSMAP, which was then verified on SeaAD. After applying the same labeling approach as for ROSMAP, we are left with 33 AD samples and 7 healthy controls (43 are labeled OTHER and excluded from determining the classification performance).

#### COMBAT

For the COMBAT dataset, the processed CITEseq data provided by the original study was used.^22^ We used the SARS-CoV-2 PCR data to split the samples between healthy (negative PCR) and diseased (positive PCR), of which there were 45, and 79 respectively.

### Baselines

We used the following four baseline models to compare the quantitative performance of scAGG:

- **Cell-type proportion classifier:** Each sample is represented by its abundance of each of the annotated cell types, normalised to 10*k* cells per sample. The resulting features (cell type proportions) are then standardised over all samples. The resulting sample representations are classified using a Lasso regression model with regularisation strength *α* = 0.03.
- **Pseudobulk classifier:** Raw gene expressions of each sample are aggregated into a single vector per sample by adding the counts per gene in all cells. Resulting representations are normalised to sum to 10*k* counts per sample and log-transformed. The classifier is an MLP with two hidden layers (hidden dims 512 and 256).
- **Cell-level classifier:** Sample-level labels are projected onto each underlying cell, assuming that all cells in a sample are affected by the disease. The classification model is a Lasso regression with *α* = 0.3. To compare these cell-level predictions with the sample-level predictions, we aggregate the results by considering the proportion of cells that were classified as AD in each sample, and pick the optimal decision boundary using the train set.
- **ScRAT**^12^ Attention-based classifier, used to represent the state-of-the-art, which was most reproducible out of all existing methods discussed in the introduction. We use the default hyperparameters; no additional tuning was performed.

### scAGG: Sample-level embedding and classification architecture

We propose scAGG (single-cell AGGregation), a three-part architecture that consists of (1) a cell-level encoder, (2) a sample pooling mechanism, and (3) a sample-level classifier. These three modules can be trained in an end-to-end fashion on the sample-level disease classification task. We first describe the abstract modules, after which we define the specific parametrizations we evaluated in this work.

We define the *cell encoder* module as a learnable mapping from the input space χ ∈ ℝ^*n×m*^, to a latent space Ƶ ∈ ℝ^*n×h*^, where *n* denotes the number of cells, *m* the number of features (genes) and *h* the latent dimensionality.

The *pooling mechanism* is defined to take a set of cell embeddings *Z* ⊂ Ƶ belonging to a single sample (donor) and return a sample-level embedding 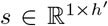. Note that the latent dimensionalities of cell- and sample-level embeddings are not necessarily the same.

Finally, we define a *classifier* module that predicts the disease state *Ŷ* ∈ ℝ^*N ×*2^ for each sample-level embedding *s*. In all variations of the model we explore, this classifier part of the architecture is the same.

### scAGG: Base parametrization

This model chooses the simplest parametrizations for all three modules of the architecture. The encoder is a two-layer neural network, with a hidden dimension of *h* = 128, interspersed by a non-linear activation function (ReLU):

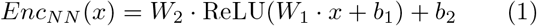

The pooling layer is a global mean pooling layer *P*_*mean*_, which averages the cell-level embeddings to a single graph-level embedding:

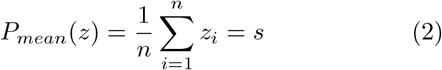

The classifier is a linear layer, followed by a softmax activation function.

### scAGG+AP: Attention-based pooling

To allow a sample pooling mechanism to prioritize specific cells when aggregating them, we incorporate an attention mechanism. We adapt the self-attention mechanism to our sample-level pooling task, using a classification token *T*_*CLS*_ ∈ ℝ^*h*^ as was used, for example, in BERT.^23^ This is a learnable token that is appended to the set of cell embeddings for each individual sample 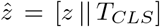, used to represent the entire sample and calculate the attention scores for each cell. Using the classification token, we can use the self-attention mechanism without any modifications. To increase model capacity, we use four attention heads, which have separate weights, and the resulting embeddings are averaged to obtain the final sample-level embedding.

### scAGG+GAT: Graph attention cell encoder

To incorporate attention into the cell encoder part of the model, rather than the sample pooling layer, we incorporate a Graph Attention Network (GAT).^24^ Here, each sample is represented as one kNN-graph, where each node represents a cell, and edges are added for each cell to its *k* = 30 nearest neighbours, according to Euclidean distance in the space spanned by the first 50 principal components of the gene expression.

Given a graph *G*, and the function *N*_*k*_(*i*) that denotes the set of the *k*-nearest neighbours of a node *i*, a GAT layer calculates an embedding for each node *i* as follows:

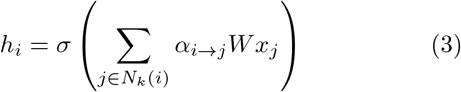

Here, *x*_*j*_ is the feature vector of neighbour *j*, and *W* is a learnable linear transformation. The attention *α*_*i*→*j*_ that is paid to *j* when calculating the embedding of *i* is calculated following the definition of GATv2, where different weights are learnt for *i* and *j*, to allow for “dynamic” attention:^25^

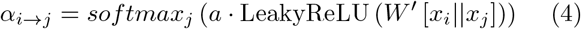

where *a* and *W* ^*′*^ are learnable transformations, and the || operator denotes vector concatenation. Note that, unlike most GAT implementations, we do not include self-loops: *i* ∉ *N*_*k*_(*j*), for better interpretability of attention scores. We use 8 attention heads and concatenate the resulting embeddings before passing them to the classifier.

The classifier and pooling layer are the same as in the base parametrisation, although we also consider a combination of the GAT encoder with the attention-based pooling in our experiments (scAGG+GAT+AP).

### Model Training & Evaluation

All models were trained and evaluated using a 5-fold cross-validation, stratified by disease class, where the same splits are used between models to allow for a fair comparison. Splits were stratified by disease class at the donor level. That is, we hold out entire donors in each test set. We used a categorical cross-entropy loss function and the model weights were optimised using Adam^26^ (*lr* = 0.001, *wd* = 0.0005 and dropout rate of 0.5 for the GAT-based models, and 0.1 otherwise. Dropout was applied after every encoder layer).

Mini-batches for graph-based model were constructed using neighbour sampling,^27^ where eight seed nodes are sampled from the different training graphs, in a weighted fashion to correct for class imbalance and difference in cell counts between samples. For each seed node, 15 random neighbours are added to the batch graph and for each neighbour, 15 additional (2-hop) neighbours are added to the batch graph. As a result, each batch has at most 8 *×* 15 *×* 15 = 1800 nodes. In the original neighbour sampling algorithm, all nodes are used once as seed nodes every epoch, but in our case, we define an epoch as a training iteration over only 200 of such batches, to allow for early stopping, because otherwise the model would overfit within a single epoch.

For the models that do not rely on a graph structure, we found it worked equally well to train on batches generated in the same way, so we use the same batching technique for all models for consistency, and discard the graph structure after sampling for the non-graph models.

When replicating our approach to SeaAD, we trained each model on the ROSMAP dataset (excluding intermediate cases like before), based on a subset of 959 genes that overlap with the 1000 HVG from SeaAD, and using the same hyperparameters as described for the other experiments.

### Polygenic Risk Scores

To calculate the polygenic risk scores (PRS) for each donor, we collected weights for known risk SNPs reported by Wightman et al..^28^ These do not include the primary AD risk SNPs, rs429358 and rs7412, that define the APOE e4/e2 alleles, as is customary for genetic AD studies. Out of the 38 reported SNPs, 35 were available in the wholegenome sequencing (WGS) data of ROSMAP. Since the WGS data reports dosages, we had to flip dosages where the opposite strand was measured. Resulting dosages were used as input to a linear regression with the aforementioned weights to obtain the PRS scores for all ROSMAP donors. For visualization purposes, the PRS scores were standardized.

### Within-individual Heterogeneity

To characterize within-individual heterogeneity in a per-celltype fashion, we considered the first principal component of the cell-level embedding space for each celltype separately, representing a celltype-specific disease axis. We then calculate a donor’s celltype-specific involvement by taking the mean over all their cells’ positions along this axis, resulting in a score per donor per celltype.

### Attention Analysis

To identify the cells prioritised by the attention mechanism, we calculated the received attention of each cell *i*, by summing over attention it received from all neighbours *j* ∈ *N* (*i*). The score is then corrected for the degree of the cell, using the square root of the degree, so that high-degree cells are penalised less harshly. Finally, we scale the overall distribution to sum to 1 again by multiplying by the average degree of the sample *k*:

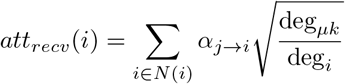

To investigate the selectivity of the attention mechanism, we analysed the entropy of the outgoing attention weights of each cell:

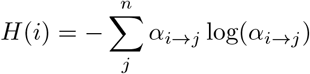

Note that the direction of the attention here is opposite to the received attention, because selectivity of the attention mechanism occurs in the outgoing attention, whereas the received attention is more indicative of the importance of a cell in the prediction task.

The attention entropy distributions are then compared with a uniform distribution, which is obtained by setting all the attention weights in a graph to 1*/n*, where *n* is the number of neighbours of the cell, and then applying the above formula again.

### Computational Resources

The full training procedure for scAGG on the ROSMAP dataset required 22 GB of RAM and took 8 : 45 minutes on a modern desktop PC using the GPU (RTX 4070 Ti). For the attention-based implementations, the memory requirement remains equal, but runtime increases to 10 : 48, 10 : 47 and 16 : 55 minutes for GAT, AP and GAT+AP respectively. Furthermore, training time scales linearly in the number of cells and genes, except when using the GAT encoder, which scales quadratically in *k*, the number of neighbors in the KNN graph. The runtime of the attention-pooling layer scales quadratically in the number of cells per batch, but this does not increase with the overall number of cells.

## Results

To address the task of classifying cell-level data with sample-level labels, we propose scAGG, a three-part architecture that consists of (1) a cell-level encoder, (2) a sample pooling mechanism, and (3) a sample-level classifier (Fig 1). In its base form, which we refer to as scAGG, the cell encoder and sample classifier are both simple neural networks, and the sample pooling mechanism is a global mean pooling function that aggregates cell-level embeddings into a sample-level embedding. This sample-level embedding is then used to classify the sample as a whole (see Methods).

**Figure 1:**
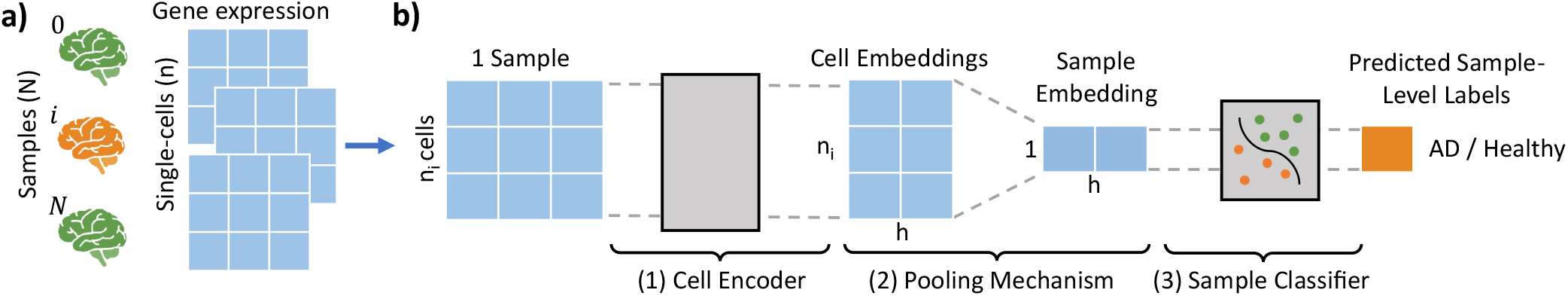
Illustration of the proposed scAGG model: (a) For *N* samples (brain donors) we have a total of *n* cells, each of which have *n*_*i*_ cells (b) The cell encoder module (1) produces *n*_*i*_ cell embeddings that are aggregated by the pooling mechanism (2) into a single embedding per sample. The resulting sample embedding is then classified according to disease status using a classifier module (3).

### Datasets

To train and evaluate, we used ROSMAP,^17^ a single-nucleus RNAseq dataset consisting of dorsolateral prefrontal cortex (DLPFC) samples from patients with Alzheimer’s disease and healthy controls of various stages of AD, providing a comprehensive view of the disease and its progression (Fig S1). After processing 339 donors (2, 037, 478 cells) remained, for which the 1, 000 most highly variable genes were selected. Following the approach proposed in (Wang 2022)^20^ we identified the most extreme cases of AD (78) and the least cognitively impaired controls (61), leaving 200 intermediate donors (see Methods). We use only the extreme cases for training and performance evaluation, so the model learns to separate the most extreme ends of the disease spectrum. In subsequent analyses, we also included intermediate cases.

To further assess scAGG’s capability to generalize across datasets, we used the Seattle Alzheimer’s Disease Brain Cell Atlas (SeaAD)^21^ as a replication dataset. Like ROSMAP, this is a snRNAseq dataset of donors from various stages of AD. We only used the DLPFC data, as this is also what we trained on in ROSMAP. After selecting the least cognitively impaired and more extreme AD cases, 40 samples remained.

We also evaluated scAGG in the context of another disease, SARS-CoV-2. For this we used the COMBAT dataset,^22^ a single-cell CITEseq dataset, containing 124 peripheral blood mononuclear cell (PBMC) samples with different severities of the disease, and healthy controls. The target labels for a binary classification task were derived from the polymerase chain reaction (PCR) test data.

### Pooling of cell embeddings for sample-level classification of extreme AD cases

To quantitatively assess the performance of scAGG, we consider the task of classifying only the most extreme cases of AD and CT, as these are expected to be the most separable cases on the disease continuum. We compare our performance with three baselines: a cell-type proportion classifier (CT Prop), a pseudobulk model, and a cell-level classifier model that does not rely on pooling together the cell embeddings (see Methods).

We find that scAGG outperforms the CT Prop and Cell-level baselines on all metrics (Fig 2), whereas it performs on par with the pseudobulk model, with a slightly higher mean F1 and AUC, although not statistically significant. However, we note that this pseudobulk approach discards any opportunity to uncover the cellular heterogeneity captured by the single-cell data. We observed a high standard deviation between the different folds of the cross-validation procedure used for scAGG, which is primarily due to sensitivity to the data split per fold, as we show the model is more stable over repeats of the same fold

**Figure 2:**
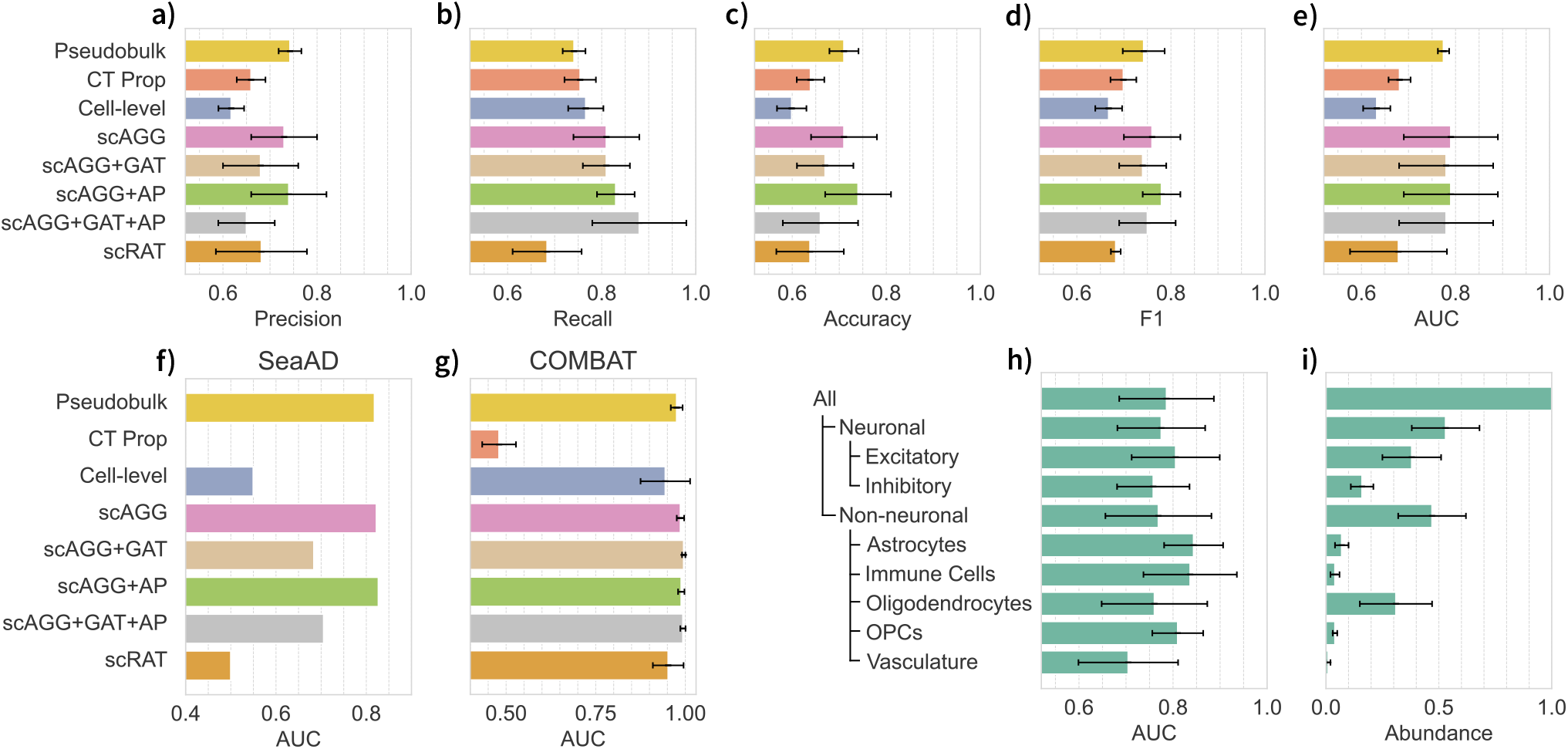
Quantitative performance analysis: a-e) Baseline models and variations of the proposed scAGG architectures are compared for the 1000 most highly variable genes; f) Performance when trained on ROSMAP and evaluated on SeaAD; g) Performance when trained and evaluated on the COMBAT dataset; h) Classification performance for several cell types; i) Relative abundance per cell type. Error bars represent the standard deviation over 5 folds of cross-validation, except for panel i where the error bars denote the standard deviation over all samples. Raw numbers can be found in Tab S1 and S2.

(Fig S2). The performance of the other baselines comes closest to the performance of scAGG on the recall metric, which we attribute to the baseline models predicting the AD class more frequently, since this is the majority class. We also compared it with scRAT^12^ (default parameters), representing the state of the art and show that scAGG has a higher mean performance on all metrics, although statistical significance could not be established due to the high variability of both methods due to the low number of samples left when conducting nested cross-validation.

To assess the effect of the chosen input set of genes, we also consider performance on the top 5, 000 HVGs (Fig S3) and we find that the performance of scAGG does not improve further. Only the performance of the pseudobulk baseline model does improve, but the AUC of scAGG is still on par. We also evaluated the performance on a set of 979 genes known to be differentially expressed in AD [10] and show this does not outperform the 1, 000 HVGs (Fig S4). The overlap between this set of known AD genes and our selected set of genes is 48 and 230 for 1, 000 and 5, 000 HVGs respectively.

We further assess the generalizability of scAGG by training without cross-validation on ROSMAP and evaluating the performance on an external replication Alzheimer’s disease dataset, SeaAD (see Methods). The CT Prop baseline was not evaluated in this setting, as we did not have the aligned cell type annotations for both datasets. When comparing the AUC (to account the for the high class imbalance, i.e. 8 AD vs. 32 control samples), we find scAGG and scAGG+AP perform on par with the pseudobulk classifier but outperform all other methods (Fig 2f).

We also consider the classification performance on a SARS-CoV-2 dataset, COMBAT (see Methods), and find that all model versions of scAGG perform equally well, together with the pseudobulk model, but scRAT and the cell-level model perform marginally worse (Fig 2g).

### Sample-level embeddings show association between AD classifiability and pathological stage

After training the proposed architecture (Fig 1), we obtain a single embedding for each individual that aggregates all single-cell data of this sample, directly optimised for the classification task. To illustrate how the embeddings of the samples clearly separate disease states, we show the distribution of disease classes along the first principal component (PC) of the sample-level embedding space.

Using the model that was trained on the extreme cases only, we projected the held-out intermediate samples onto this spectrum. These intermediate samples are mapped between the disease extremes (Fig 3b), illustrating that the sample embeddings learnt by our model capture a healthy-to-AD disease gradient, while only trained on a binary classification task. Moreover, we show that this ordering aligns with the disease severity of the samples, as quantified by the Braak stage and CERAD score, a pathological marker and a clinical score respectively (Fig 3c+d). This indicates that a sample’s position on this learnt spectrum can be interpreted as a measure of disease severity.

**Figure 3:**
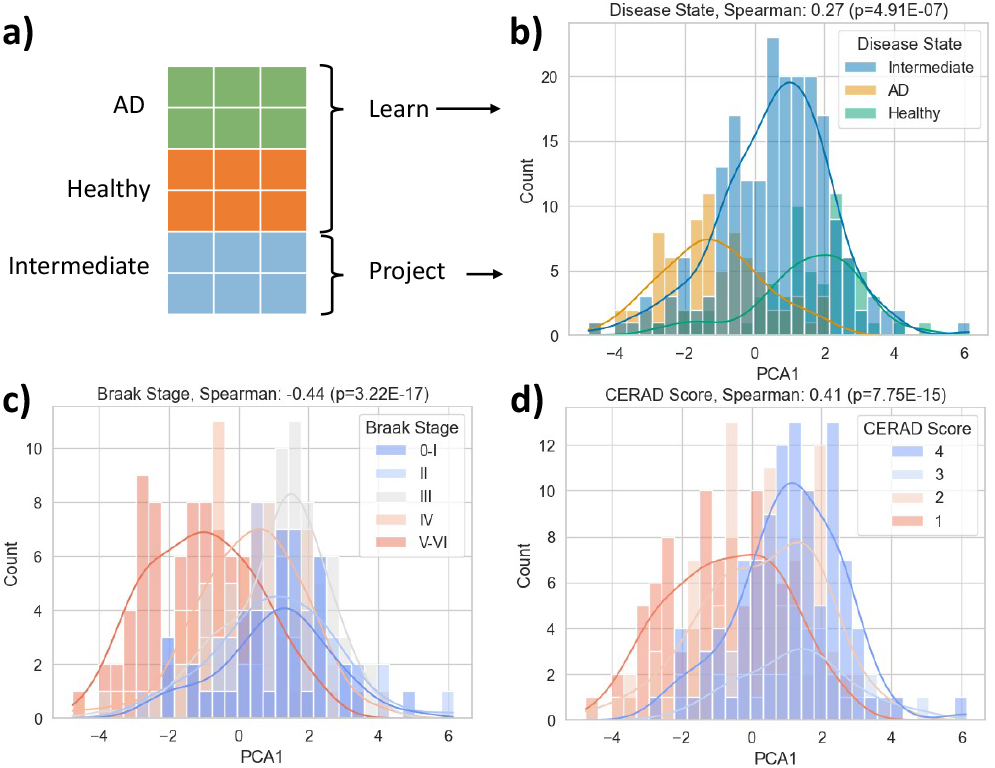
Visualization of sample-level embeddings, showing the distribution of classes along the first principal component, coloured by: (b) disease status of extreme cases, including intermediate cases projected onto latent space (as illustrated in panel a), (c&d) Braak stage and CERAD scores of all samples, including intermediates.

As AD status is confounded by both sex and age in the dataset (Fig S1), we assessed the effects of these variable in the resulting embedded space. We found that sex and AD status are both significantly associated with the first PC (*p* = 0.000, and *p* = 0.001 resp.), but AD status has an over 2.5*×* larger coefficient (Tab S3).

### Cell type-specific performance differences

The aforementioned approaches all rely on the use of all cells in a sample to make a prediction, but these cells belong to different cell types, which may not all be equally informative for the prediction of AD. Therefore, we investigated the classification performance of several major cell types by training and evaluating the model only on cells of that type. We find that there is little difference in performance when considering all neuronal vs. all non-neuronal cell types, which in have similar relative abundances in total. Once we consider individual cell types, we find that, for neuronal cell types, excitatory neurons are more accurately classified, but this could be due to their higher abundance. For non-neuronal cells, we find that astrocytes, immune cells (including microglial cells), and OPCs can be classified exceptionally well considering their far lower abundance compared to oligodendrocytes, which make up the majority of the non-neuronal cells, while being far less accurately classified (Fig 2h).

When we further consider the classification performance at a higher cell-type resolution, we find that GRM3-expressing astrocytes, in particular, are well classified, together with all 5 different types of RORB-expressing excitatory neurons (Fig S5).

In general, this indicates that some cell types are more predictive of AD than others, which motivates the use of a model that can prioritise specific cells when making a prediction.

### Attention-based sample pooling mechanism does not outperform naive global pooling

Having shown that some cell types are more predictive of AD than others, we propose to incorporate an attention mechanism^29^ that allows the model to prioritise specific cells when making a prediction, to make better predictions with only the most informative cells. First, we consider an attention-based sample pooling mechanism (scAGG+AP). Instead of using a simple global sample pooling that averages all cell embeddings, we use an attention mechanism that learns a scoring mechanism per cell that determines its weight in the global embedding (see Methods).

We find that the attention-based sample pooling mechanism does not improve the performance of the model (Fig 2). Only when increasing the number of HVGs to 5, 000 do we see a slight improvement in performance compared to the naive sample pooling mechanism in terms of accuracy, but the F1 and AUC are similar (Fig S3). This indicates that the attention-based sample pooling mechanism overfits to the majority class (AD), improving the total accuracy, but not when corrected for class imbalance.

### Graph-attention based cell encoder module does not improve performance

The global application of attention as presented above scales quadratically with respect to the number of cells and can become computationally expensive, since the scRNAseq samples may consist of 10, 000s of cells. In addition, it embeds each cell independently and does not consider the relationships between cells, while it has been shown that incorporating information from similar cells can be valuable.^5,6,8^

To address this, we investigated another approach in which we employ a Graph Attention Network (GAT).^24^ In this approach, each sample is represented by a cell graph, and the GAT learns to incorporate information from neighbouring cells into the cell embeddings. An attention mechanism is incorporated that learns weights for the neighbours of a cell, which can later be interpreted (see Methods).

We evaluated the quantitative performance of the GAT-based encoder module combined with both the global mean pooling approach (scAGG+GAT) and the attention-based sample pooling (scAGG+GAT+AP). We find that the GAT-based encoder does not improve the performance of scAGG (Fig 2). On most metrics, the performance is either slightly lower, or on par with scAGG. Only the recall metric is higher for the GAT-based encoder with attention-based sample pooling, which we attribute to the model overfitting to the majority class.

### Cell-level embeddings show separability between disease classes

In addition to considering the sample-level embeddings, we can also consider the cell-level embeddings before they are pooled together into a sample-level embedding. We compared a UMAP^30^ reduction of the input space with the cell embeddings produced by scAGG and the GAT-based model (Fig 4).

**Figure 4:**
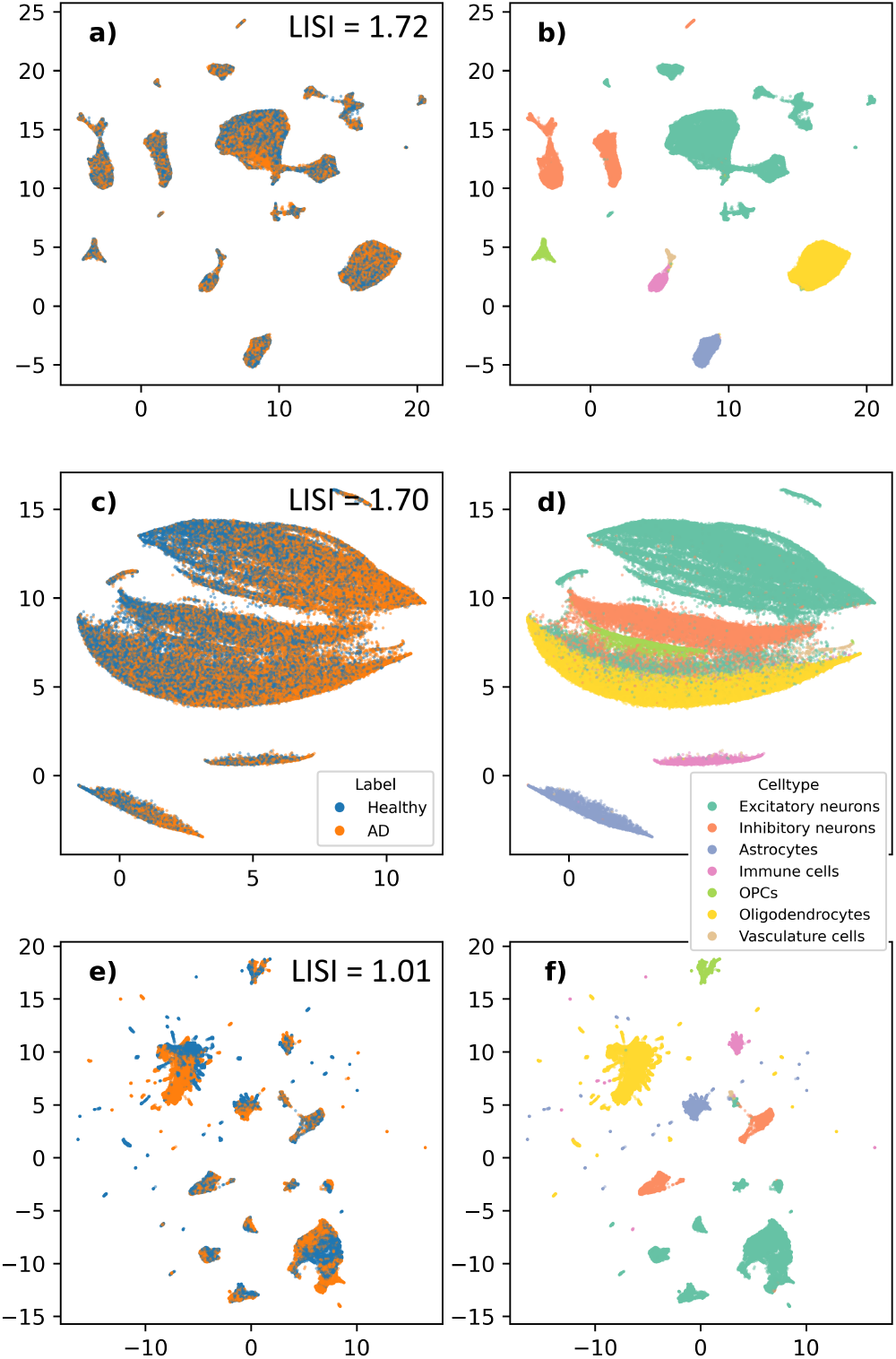
2D UMAP visualization of the embeddings at the single-cell level, coloured by disease status (left column), and cell type (right column), showing the differences in representations from the input space (a & b), the baseline model (c & d), and the GAT model (e & f). The cell embeddings learned by the attention-based pooling model is very similar to the scAGG embeddings and therefore omitted.

Starting with the input space, we see that there is a clear separation between the different cell types (Fig 4b), but little apparent separation between the disease classes (Fig 4a). When we compare this with the embeddings produced by scAGG, we observe that cells are arranged along a healthy-to-AD gradient (Fig 4c). The GAT embeddings appear to be noisier than the other two representations but still show separation between disease classes within celltype clusters (Fig 4e+f).

Investigation of UMAPs per cell class (Fig S6) further shows that the scAGG cell-embedding space is primarily organized by AD association of cells and less by celltype-specific patterns. These patterns also seem to correspond partially with polygenetic genetic risk (see Methods), although we do not detect distinct subpopulations with particularly high or low genetic risk. At this scale, the input space also shows clearer separation between disease states, but in a less organized fashion.

To quantify disease class separability, we compared the LISI scores for both types of embeddings and input space, which quantify the number of classes that occur in a cell’s neighborhood. For perfect separation, we expect *LISI* = 1 and for a homogeneous mixture we expect *LISI* = 2. For the input space, we find a mean LISI score of 1.72, compared to scores of 1.70 and 1.01 for the scAGG and scAGG+GAT models, respectively. For scAGG, this corresponds to the visual observation that the majority of the space is mixed, and only at the extremes are more homogenous. The much lower LISI score for scAGG+GAT is also represented in the per-celltype UMAPs, where we observe that cells are arranged in clusters per donor (exemplified by clusters having the same genetic risk score) (Fig S6). This indicates scAGG+GAT overfits to donor-specific patterns, which may explain why it does not outperform scAGG on the classification task, despite its increased latent separability.

### Cell-level embedded space associated with several ion response pathways

The UMAP embeddings of the cell-level embeddings learnt by scAGG (Fig 4c) align with a healthy-to-AD spectrum. This is even more pronounced when we consider the first principal component of cell-level embeddings for a single cell type, such as astrocytes (Fig 5a). Here we see that the first PC of the cell embedding seems to order cells along a healthy-to-AD spectrum. To characterise the gene expression patterns that are associated with this spectrum, we calculated the Spearman correlation between the gene expression of each cell and the corresponding value of the first PC. Most genes show a correlation close to zero, but some genes show a strong correlation (Fig 5b).

**Figure 5:**
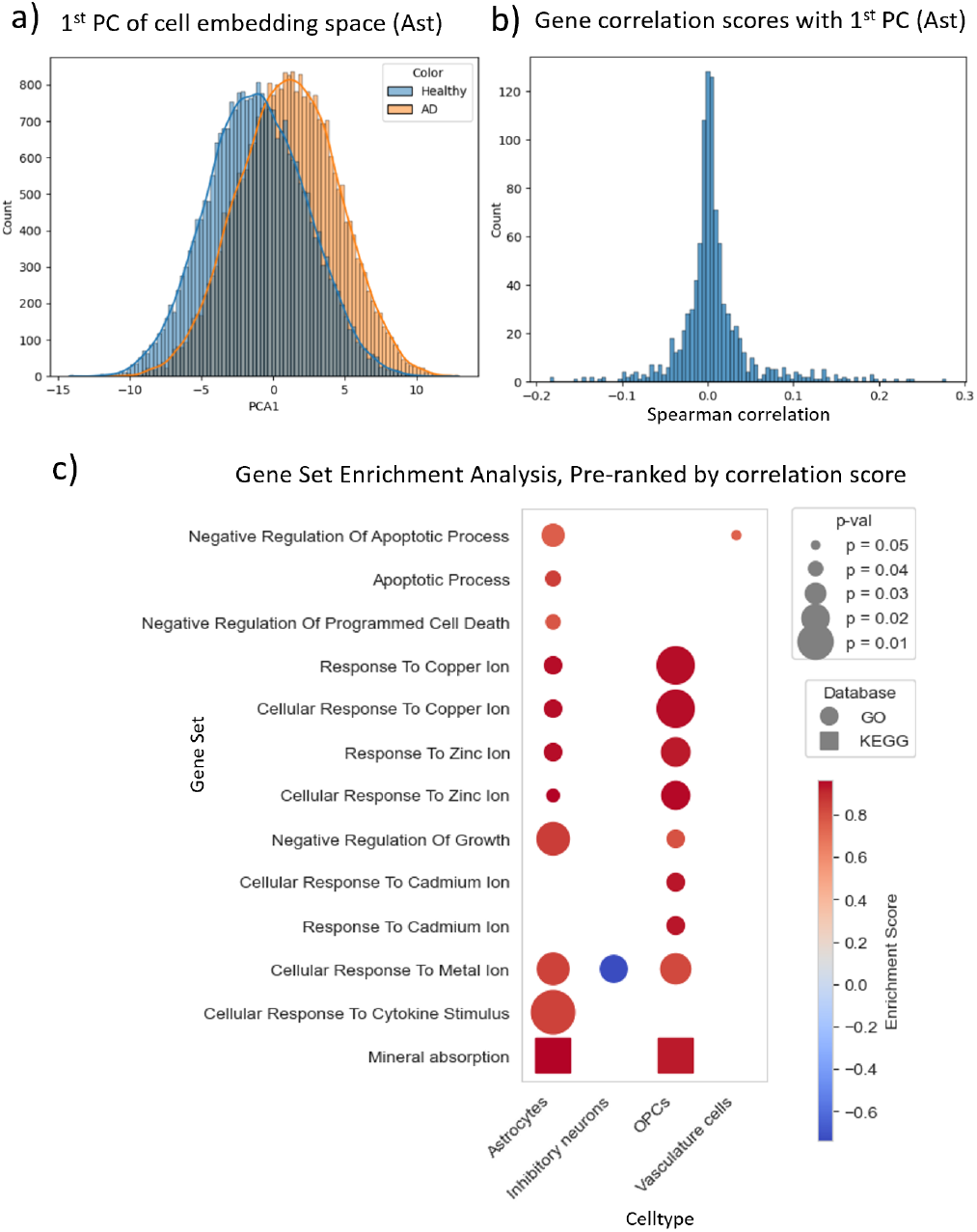
Analysis of the learned cell embedding space of scAGG. (a) The first principal component all astrocytes, coloured by disease class. (b) Distribution of the correlation scores for each gene, obtain using the spearman correlation between the gene’s expression in each cell, and the corresponding cell’s 1st PC value. (c) Gene set enrichment analysis per cell type, using the obtained correlation scores to pre-rank the genes.

We then use the absolute values of these correlation scores to rank genes and perform a gene set enrichment analysis (GSEA) for each cell type separately (Fig 5c). We find several significantly enriched gene sets in both the `KEGG (2021) Human Pathways` and `GO Biological Process (2023)` databases for 4 of the 7 major cell types. In particular, we find pathways related to metal ion signalling, regulation of cell growth, and programmed cell death. Furthermore, we find that there is substantial overlap between the enriched gene sets of the different cell types, with 8 out of the 13 gene sets being significantly enriched in at least two cell types.

### Within-individual heterogeneity

The fact that the cell-level embedding space of scAGG arranges the cells according to disease severity allows us to assess a donor’s disease state at the cell level and for each cell type separately. In a heterogeneous disease such as AD, it may be the case that some donor’s cells are spread over the entire spectrum, while they are highly concentrated at one of the extremes for another (Fig S7). The same differences are possible between different cell types within the same sample. To identify such cases of within-sample heterogeneity, we use the learned scAGG embeddings to characterize celltype-specific involvement (see Methods). We find that these scores largely correlate between the different cell types but identify a few cases where AD-diagnosed donors show less severe astrocyte scores. Among the cognitively healthy controls, there also seems to be a divide between mostly healthy donors and donors where mainly the glial cells are still on the least-impaired end of the spectrum, but the neuronal cells’ scores are less pronounced (Fig 6).

**Figure 6:**
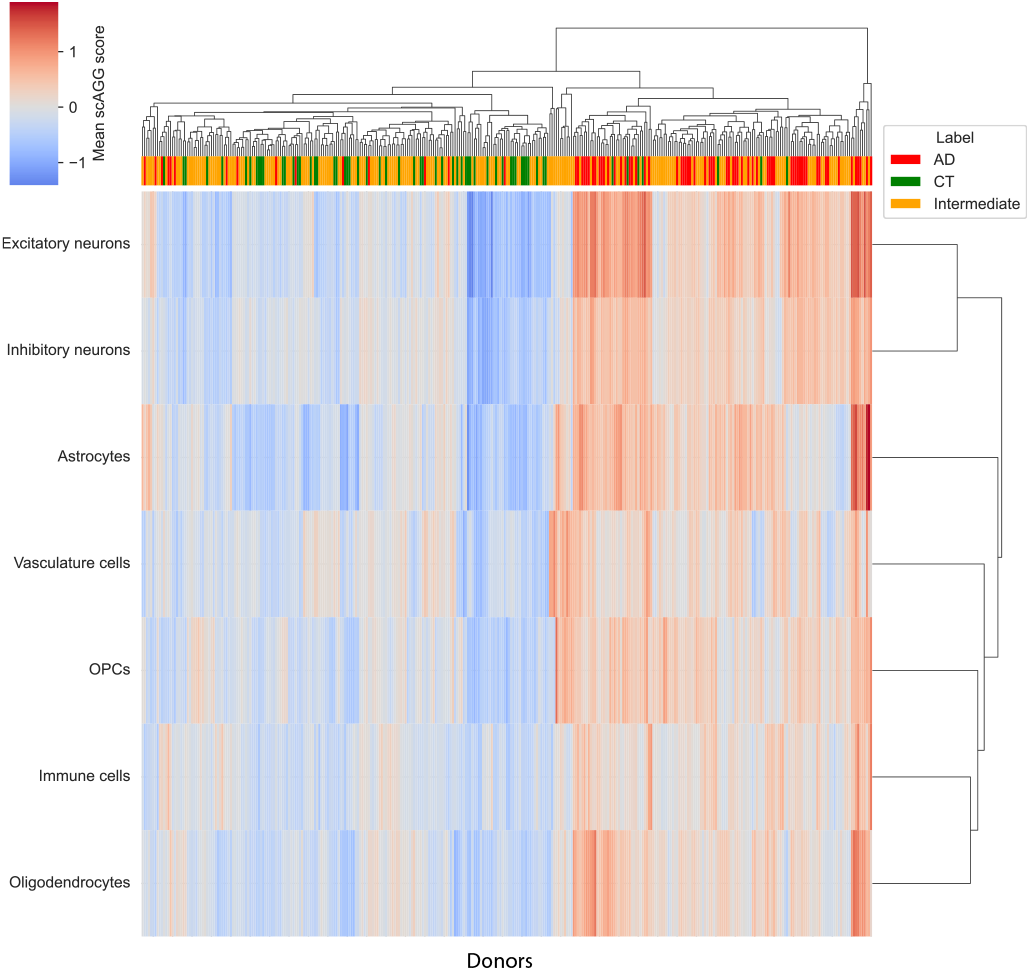
Celltype-specific disease association scores of all donors, per major cell type. Rows and columns are clustered based on similarity. The same analysis was also performed for all minor cell types, the results of which are shown in Fig S8.

### Entropy analysis of attention scores underscores the limited value in interpretation of attention scores

We found that the attention-based models do not out-perform simpler parametrisations of scAGG, raising the question whether the interpretation of the attention scores is meaningful from a biological point of view. The distribution of received attention scores per cell type shows the same average across major cell types, with differences in spread (Fig 7a) are largely explained by cell type abundance (Fig 7b).

**Figure 7:**
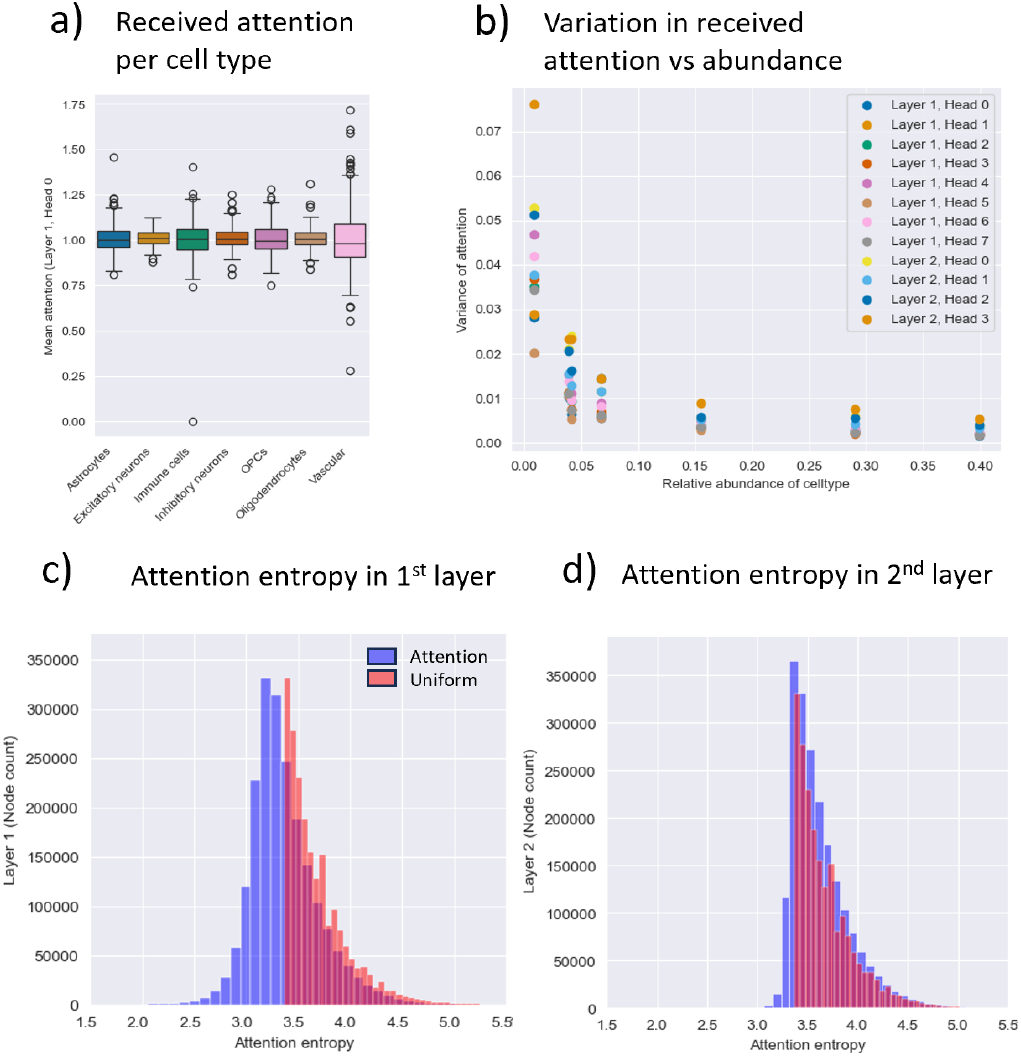
Analysis of the attention scores. (a) Comparison of distributions of received attention per major cell type. (b) Variation observed in the received attention per cell type, compared with the relative abundance of the cell type. (c+d) Attention entropy in the 1st head of the 1st and 2nd GAT layer respectively. Only the first head of each layer is shown, as the distributions are highly similar between different heads in the same layer. Blue denotes the distribution of entropy in the attention given by each cell, and red is the uniform distribution, illustrating what the entropy would look like if all neighbours were paid the same amount of attention. A single neighbour receiving all the attention would result into an entropy of 0 for this cell. Uniform attention is at least *log*(*k* = 30) = 3.5, since our graphs are built with *k* = 30.

The entropy in the paid attention scores, measuring the spread of the attention distribution, closely matches the uniform attention distribution that characterises the maximum possible entropy, in both layers of the scAGG+GAT model (Fig 7c). In the second layer, the distributions are nearly identical (Fig 7d). From this we conclude that the model spreads attention evenly rather than selectively, which explains why attention-based models do not outperform simpler alternatives.

## Discussion

We introduced scAGG, a three-part architecture for the purpose of sample-level classification of single-nucleus data, and showed that using a simple parametrisation of this architecture using neural networks already performs well. Through analysis of the learnt sample-level embeddings, we find that the latent space aligns well with two pathological markers of AD, the Braak stage and CERAD score. This indicates that the learnt space spans a spectrum of disease severity, which can be used in future work to identify cell subtypes associated with the disease and potential pathological targets.

Through evaluation of classification performance on isolated cell types, we found that the gene expression patterns of inhibitory neurons and oligodendrocytes were the least predictive of AD, despite their high relative abundance. However, this does not mean that they are not involved in the disease, because we still get reasonable classification performance from just these cell types. This is corroborated by findings of earlier work that show that all major cell types in the brain are transcriptionally affected by the disease, albeit in different ways.^**9–**11,31,32^ At the sub-celltype level, we further identified GRM3-expressing astrocytes and RORB-expressing excitatory neurons to show especially high classification performance, which are respectively known to be associated to sex-dimorphic effects on memory,^33^ and neurons that are lost early in the disease course.^34^ Furthermore, we find that performance is highest when considering just astrocytes or immune cells, despite their much lower abundance. The fact that the performance for these cell types in isolation is better than the overall performance indicates that they are overshadowed by the more abundant cell types. This led us to believe that the incorporation of an attention mechanism could further improve performance, as this would be able to prioritize the more informative cell types. We considered two attention-based parametrisations of scAGG but found that the performance did not increase as we had expected it to, based on the earlier success of attention mechanisms in similar applications. We hypothesise that, in the case of scAGG+GAT, this is because the attention mechanism can only differentiate between neighbours of a cell, which are by definition of the KNN graph highly similar in terms of gene expression and therefore generally not very informative. This warrants the use of deeper GNNs, but these will be difficult to train with the available data. With more data available to train, graph attention could improve the performance. Since the attention pooling model scAGG+AP applies attention to all cells in a sample, it should not suffer from the same problem. Yet attention did not improve performance. We speculate that this is due to the highly abundant cell types being prioritized by the pooling mechanism because they are more readily available over all samples. Future work could explore pooling mechanisms that incorporate prior information on cell type abundances to overcome this. Another limitation of scAGG is its reduction of AD classification to a 1-dimensional binary classification task, while in reality AD is a multi-faceted and highly heterogeneous disease. Future work should investigate the possibility of extending scAGG to continuous and multi-task settings, such as directly predicting Braak and CERAD scores, to reflect this complexity, but this will introduce new challenges regarding the interpretability of learned patterns. We also observed some association between the learned embedded space and sex. While still over 2.5 *×* less apparent than the AD signal (Tab S3, this deserves deeper investigation on a larger dataset.

Genes whose expression was correlated with the learned AD severity scores were found to be significantly enriched in several AD-associated terms. These terms included the regulation of apoptosis and programmed cell death, which is known to be triggered by A*β* and tau deposition in AD.^35,36^ We also found enrichment in cytokine signaling, where it is known that immune cells such as astrocytes and microglia in the brain release inflammatory proteins (cytokines) as part of an immune response to the presence of A*β* plaques and tau tangles.^37,38^ Moreover, we found enrichment in multiple terms related to metal-ion responses, with copper, zinc, and cadmium in particular. Existing literature has already established the involvement of metal ions in AD,^39^ where copper and zinc dyshomeostasis has been shown to lead to the accumulation of plaques of A*β* plaques,^40–43^ and the exposure to even moderate amounts of heavy metals, and in particular cadmium, has been shown to aggravate AD progression.^44–46^ While these are not novel results, they do show that the classification is performed using genes that are known to be involved in AD. This supports the biological soundness of our approach.

Our approach for obtaining cell- and donor-level AD severity scores opens new avenues for understanding complex diseases on multiple scales. By comparing donor-level phenotypes with the cell-level severity scores, we have illustrated how cases of within-individual heterogeneity can be identified. In future work, this could be used to evaluate the presence of healthy cells in diseased individuals, which could reveal protective mechanisms at the cellular level that may inspire new therapeutic strategies. Additionally, it would be interesting to further explore the scAGG cell-embedding space, for example by overlaying it with somatic changes or environmental differences to investigate how these correlate with the AD heterogeneity at the cell level. As single-cell RNA sequencing is being applied to increasingly large cohorts, our method provides a systematic way to begin to close the gap between individual phenotypes and cellular measurements, potentially accelerating the development of celltype-specific or even personalized therapeutic interventions.

## Author contributions statement

All authors worked on conceiving the method and experiments, T.V. performed the experiments and wrote the manuscript, which was reviewed by G.B., A.M. and M.R.

## Acknowledgments

This research was supported by an NWO Gravitation project: BRAINSCAPES: A Roadmap from Neurogenetics to Neurobiology (NWO: 024.004.012).

The results published here are in whole or in part based on data obtained from the AD Knowledge Portal.

Study data were generated from postmortem brain tissue provided by the Religious Orders Study and Rush Memory and Aging Project (ROSMAP) cohort at Rush Alzheimer’s Disease Center, Rush University Medical Center, Chicago. This work was funded by NIH grants U01AG061356 (De Jager/Bennett), RF1AG057473 (De Jager/Bennett), and U01AG046152 (De Jager/Bennett) as part of the AMP-AD consortium, as well as NIH grants R01AG066831 (Menon) and U01AG072572 (De Jager/St George-Hyslop).

Research reported in this work was partially or completely facilitated by computational resources and support of the Delft AI Cluster (DAIC) at TU Delft (RRID: SCR 025091), but remains the sole responsibility of the authors, not the DAIC team.

## Supplementary Material

**Figure S1:**
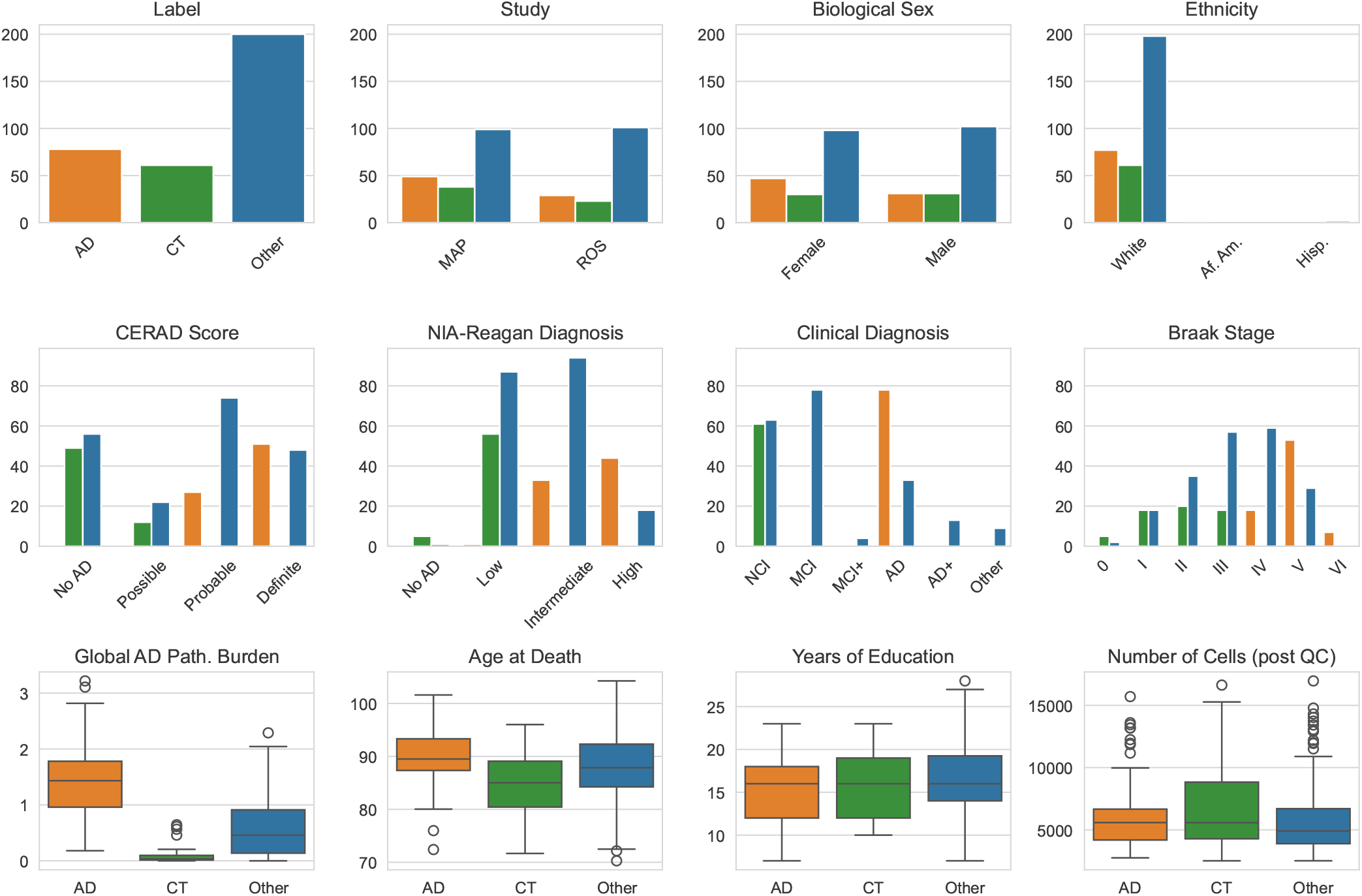
Several demographics of the ROSMAP dataset after processing and selecting the extremes of the disease spectrum, and coloured by the resulting labelling (see the Label panel for colour mapping). In the ethnicity panel, the abbreviations “Af.Am.” and “Hisp.” refer to the “Black or African American” and “Spanish/Hispanic/Latino” values in the ROSMAP metadata. In the clinical diagnosis panel, NCI=“No Cognitive Impairment”, MCI=“Mild Cognitive Impairment”, and classes with a + denote at least one additional diagnosis of another form of dementia, such as Parkinson’s disease. Braak stages increase in severity, so a higher stage is associated with higher pathology. Likewise for the global AD pathological burden.

**Figure S2:**
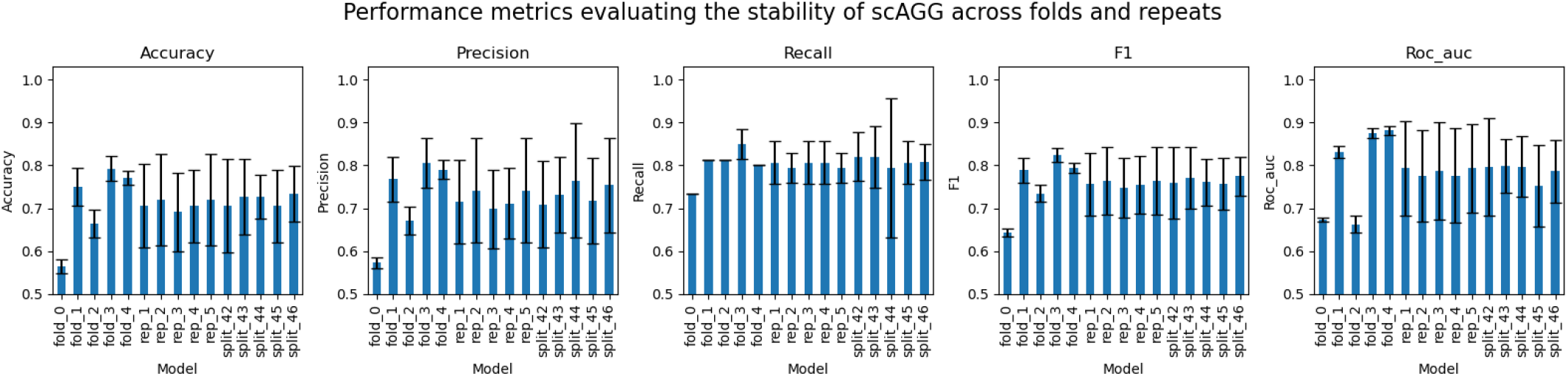
Assessment of the variability in performance of scAGG on the ROSMAP dataset with 1, 000 HVGs. We show the three different sources of variability. Bars labeled “fold” show the mean and standard deviation over 5 repeats of the same fold within a 5-fold CV experiment. Bars labeled “rep” denote repeats of that same CV experiment, using the same splits. Bars labeled “split” show the classification performance of scAGG over different stratified splits. Note that the 5 folds are part of rep 1, and the seed 42 was used for all reps. As such, rep 1 is equal to the mean of the 5 folds shown here, and split 42 should be equal to all 5 reps.

**Figure S3:**
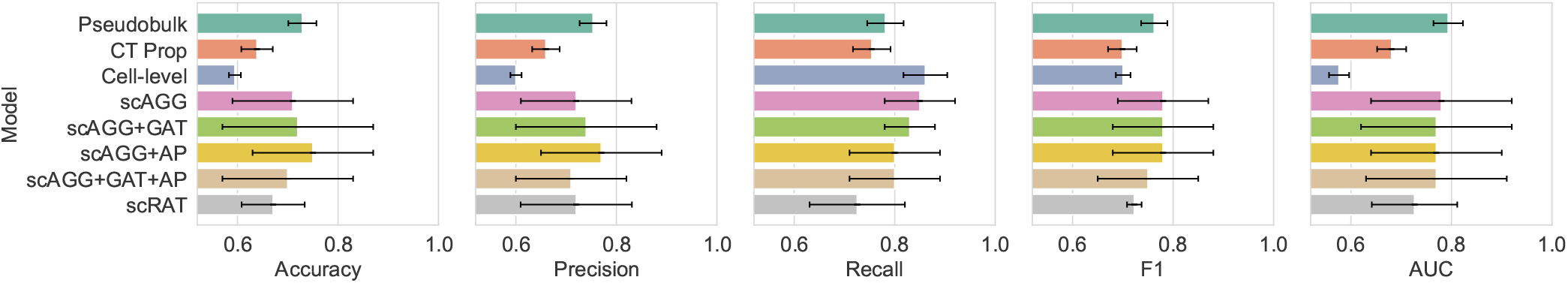
Quantitative performance analysis of baseline models and variations of the proposed scAGG architecture are compared for the 5000 most highly variable genes.

**Figure S4:**
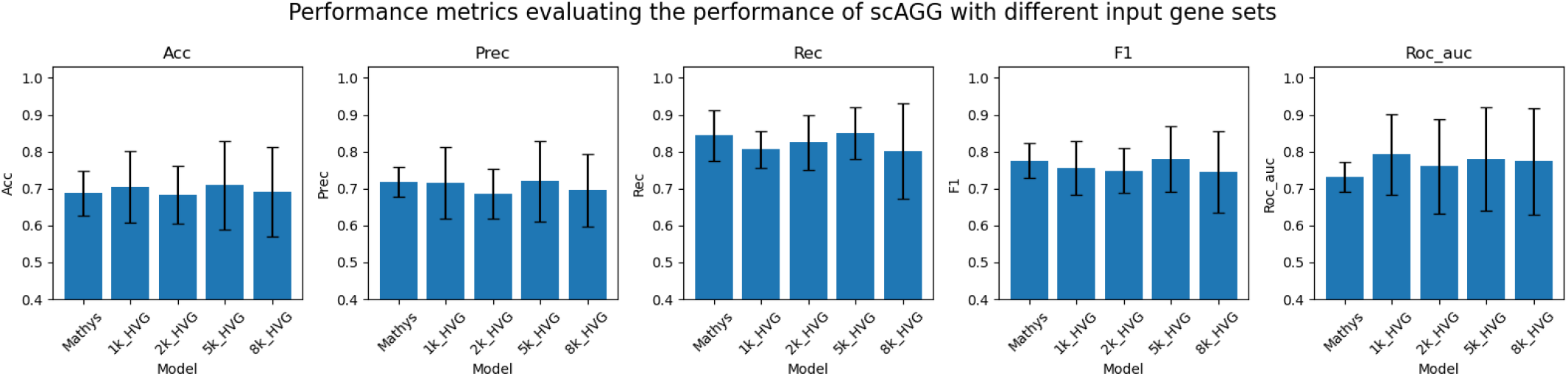
Classification performance of scAGG over different sets of input genes. Mathys denotes a set of 979 genes, which is a subset of the differentially expressed genes reported by Mathys et al.[10] that was also present in the ROSMAP dataset. The different HVG sets correspond to the output of using different values for the `n_top_genes` parameter of scanpy’s `highly_variable_genes` function. Out of these, the 1, 000 HVGs set was used throughout the paper, and performance for 5, 000 HVGs was also reported in Fig S3.

**Figure S5:**
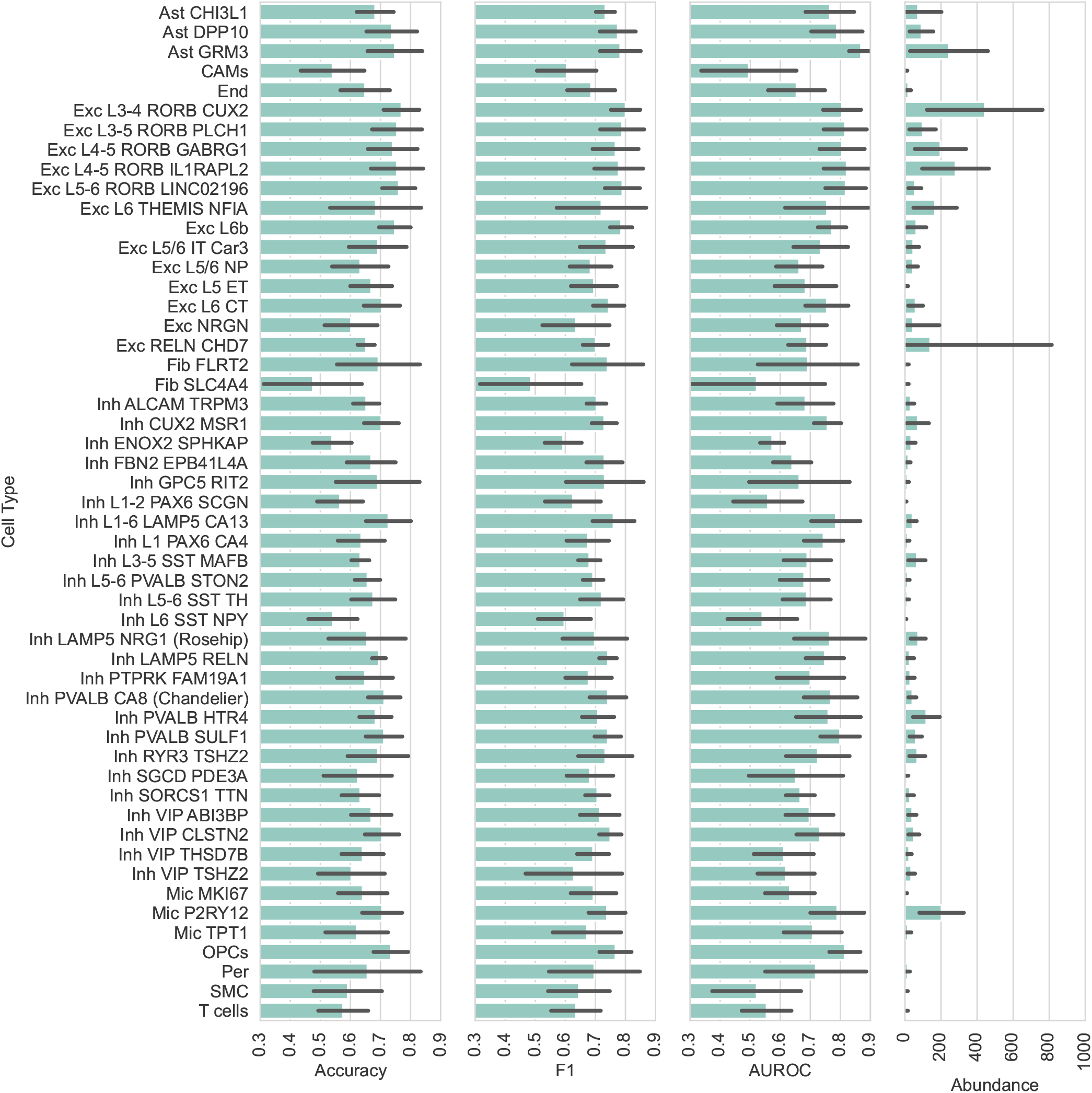
Quantitative performance analysis of scAGG on all minor cell types. The results for the major celltypes are shown in Fig 2h+i

**Figure S6:**
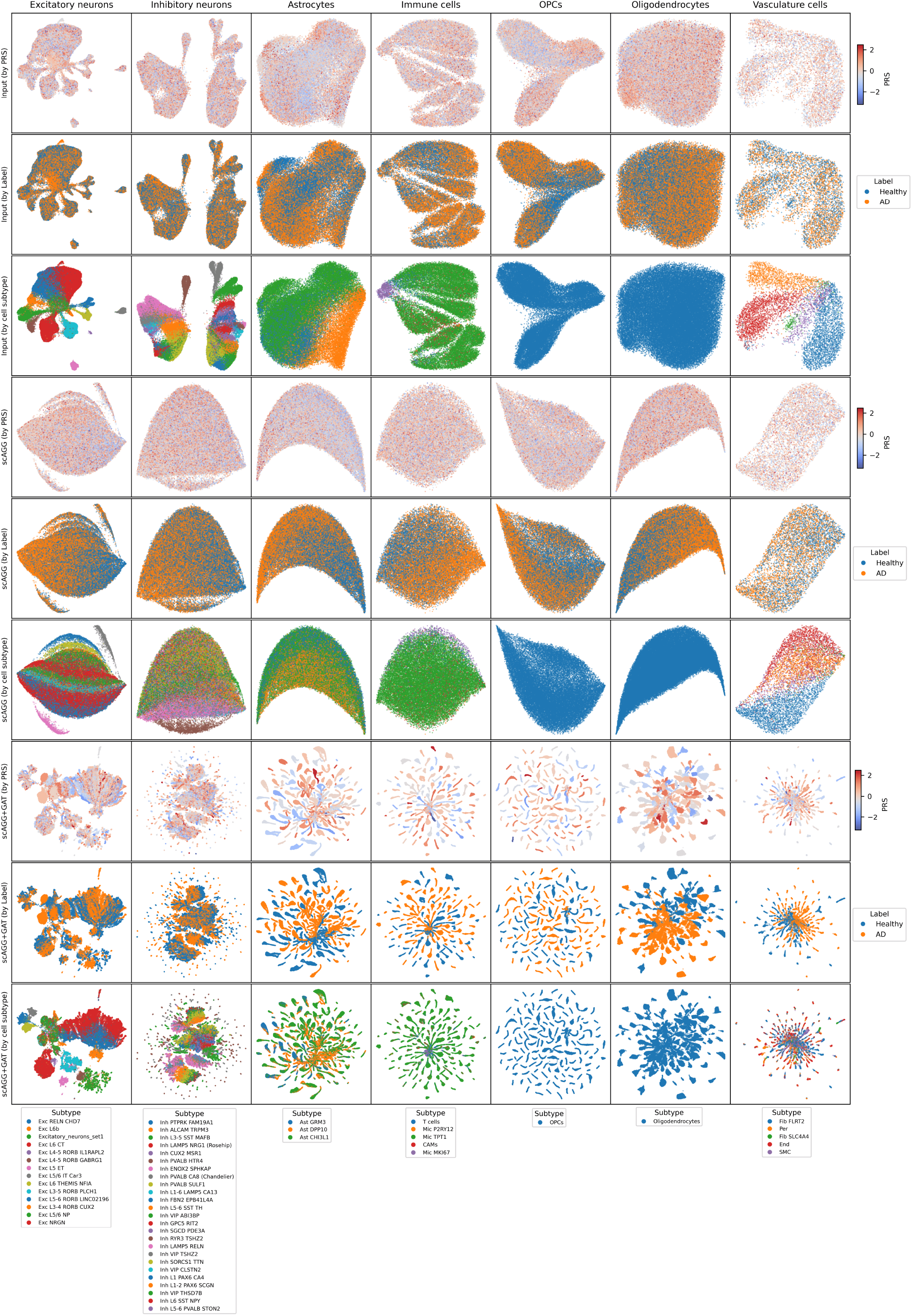
UMAP embeddings of the same three embedding spaces as Fig 4, but now per major cell type individually, coloured by polygenic risk score (every 1st out 3), disease class (every 2nd) and subtype (every 3rd). Legends for subtype are below the columns, as they differ per major cell class.

**Figure S7:**
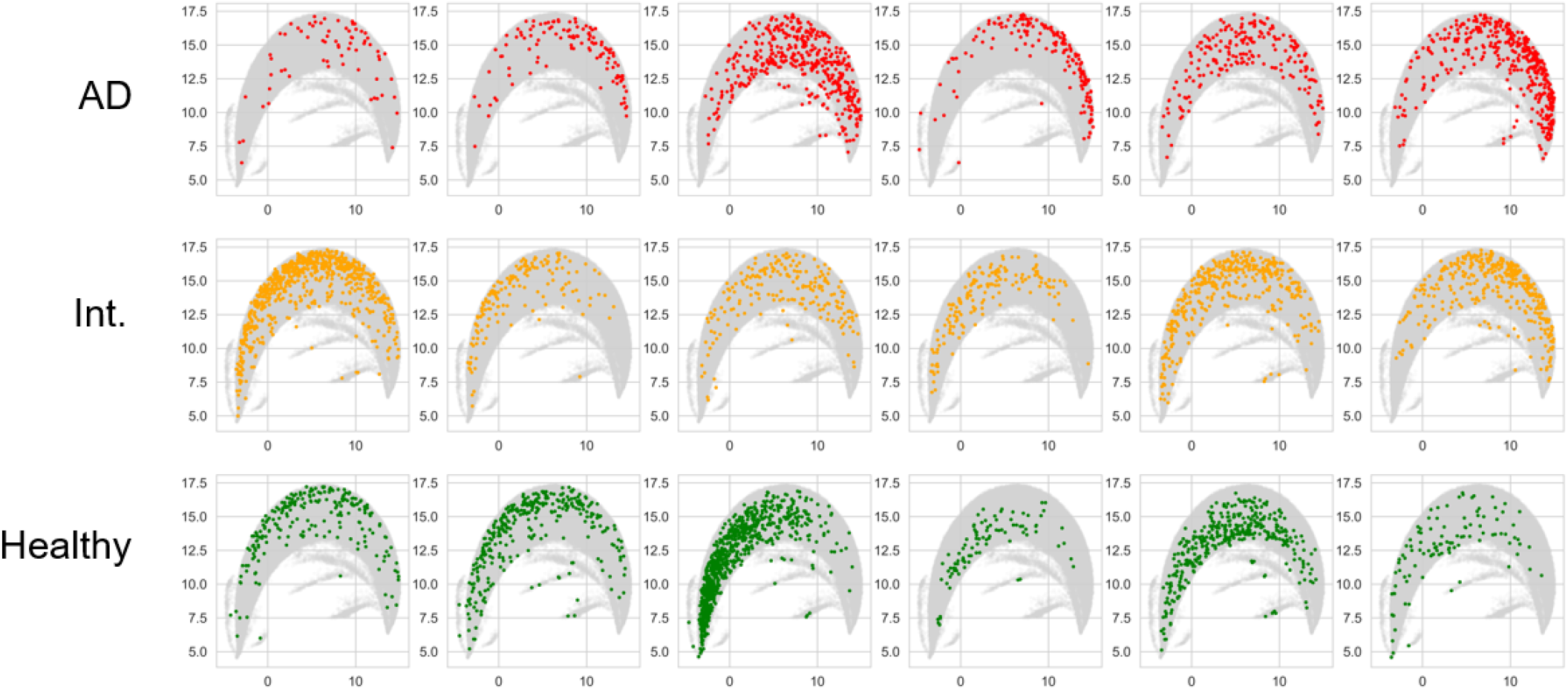
Positions of 18 arbitrary donors’ cells along the learned disease axis of astrocytes. The full UMAP of the astrocytes is shown in grey in the background, while only a single donor’s cells are colored at once.

**Figure S8:**
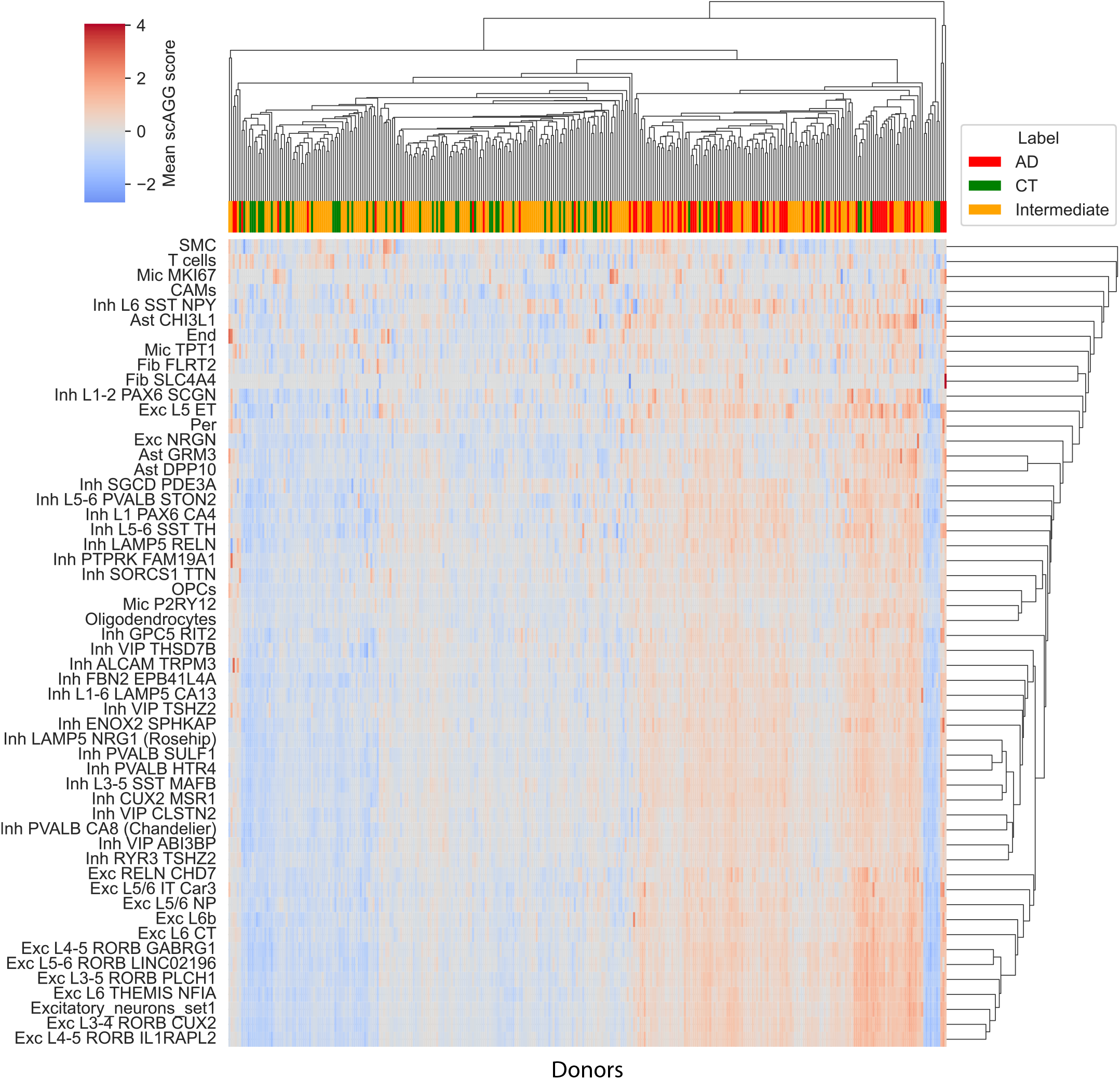
Celltype-specific disease association scores of all donors, per minor cell type. Rows and columns are clustered based on similarity. Columns are also labelled by disease state.

**Table S1:** Raw numbers for the classification performance results shown in Fig 2a-g.

| Dataset | Model | Accuracy | Precision | Recall | F1 | AUROC |
| --- | --- | --- | --- | --- | --- | --- |
| ROSMAP | Pseudobulk | 0.710 ± 0.029 | 0.743 ± 0.030 | 0.741 ± 0.033 | 0.742 ± 0.028 | 0.774 ± 0.023 |
| ROSMAP | CT Prop | 0.639 ± 0.031 | 0.660 ± 0.027 | 0.754 ± 0.037 | 0.699 ± 0.029 | 0.681 ± 0.029 |
| ROSMAP | Cell-level | 0.599 ± 0.031 | 0.618 ± 0.024 | 0.766 ± 0.024 | 0.668 ± 0.044 | 0.632 ± 0.012 |
| ROSMAP | scAGG | 0.710 ± 0.070 | 0.730 ± 0.070 | 0.810 ± 0.070 | 0.760 ± 0.060 | 0.790 ± 0.100 |
| ROSMAP | scAGG+GAT | 0.670 ± 0.070 | 0.680 ± 0.080 | 0.810 ± 0.040 | 0.740 ± 0.040 | 0.780 ± 0.100 |
| ROSMAP | scAGG+AP | 0.740 ± 0.060 | 0.740 ± 0.080 | 0.830 ± 0.050 | 0.780 ± 0.050 | 0.790 ± 0.100 |
| ROSMAP | scAGG+GAT+AP | 0.660 ± 0.080 | 0.650 ± 0.060 | 0.880 ± 0.100 | 0.750 ± 0.060 | 0.780 ± 0.100 |
| ROSMAP | scRAT | 0.638 ± 0.072 | 0.682 ± 0.097 | 0.684 ± 0.073 | 0.683 ± 0.010 | 0.679 ± 0.103 |
| COMBAT | CT Prop | 0.564 ± 0.039 | 0.654 ± 0.021 | 0.732 ± 0.082 | 0.689 ± 0.040 | 0.480 ± 0.047 |
| COMBAT | Pseudobulk | 0.914 ± 0.032 | 0.894 ± 0.039 | 0.989 ± 0.024 | 0.939 ± 0.023 | 0.976 ± 0.017 |
| COMBAT | scAGG | 0.900 ± 0.047 | 0.878 ± 0.050 | 0.989 ± 0.025 | 0.929 ± 0.033 | 0.986 ± 0.010 |
| COMBAT | scAGG+GAT | 0.979 ± 0.020 | 0.989 ± 0.024 | 0.978 ± 0.030 | 0.983 ± 0.015 | 0.995 ± 0.005 |
| COMBAT | scAGG+GAT+AP | 0.971 ± 0.016 | 0.989 ± 0.024 | 0.968 ± 0.029 | 0.978 ± 0.012 | 0.993 ± 0.007 |
| COMBAT | scAGG+AP | 0.907 ± 0.041 | 0.887 ± 0.047 | 0.989 ± 0.025 | 0.934 ± 0.028 | 0.989 ± 0.009 |
| COMBAT | scRAT | 0.912 ± 0.041 | 0.895 ± 0.068 | 0.846 ± 0.121 | 0.863 ± 0.061 | 0.952 ± 0.043 |
| COMBAT | Cell-level | 0.893 ± 0.071 | 0.884 ± 0.062 | 0.968 ± 0.047 | 0.924 ± 0.051 | 0.944 ± 0.069 |
| SeaAD | Cell-level | 0.825 | 0.842 | 0.969 | 0.901 | 0.549 |
| SeaAD | Pseudobulk | 0.300 | 1.000 | 0.151 | 0.263 | 0.818 |
| SeaAD | scRAT | 0.825 | 0.825 | 1.000 | 0.904 | 0.500 |
| SeaAD | scAGG | 0.375 | 1.000 | 0.242 | 0.390 | 0.822 |
| SeaAD | scAGG+GAT | 0.250 | 1.000 | 0.090 | 0.166 | 0.683 |
| SeaAD | scAGG+GAT+AP | 0.175 | 0.000 | 0.000 | 0.000 | 0.705 |
| SeaAD | scAGG+AP | 0.175 | 0.000 | 0.000 | 0.000 | 0.826 |

**Table S2:** Raw numbers for the celltype-specific classification performance results shown in Fig 2h+i.

| Celltype | Accuracy | Precision | Recall | F1 | AUROC | Abundance |
| --- | --- | --- | --- | --- | --- | --- |
| All | 0.720 ± 0.068 | 0.729 ± 0.073 | 0.807 ± 0.072 | 0.763 ± 0.057 | 0.786 ± 0.101 | 1.000 ± 0.000 |
| Neuronal | 0.705 ± 0.076 | 0.757 ± 0.133 | 0.758 ± 0.079 | 0.745 ± 0.050 | 0.775 ± 0.093 | 0.530 ± 0.150 |
| Excitatory | 0.734 ± 0.083 | 0.744 ± 0.095 | 0.819 ± 0.053 | 0.777 ± 0.066 | 0.806 ± 0.094 | 0.380 ± 0.130 |
| Inhibitory | 0.670 ± 0.084 | 0.688 ± 0.079 | 0.768 ± 0.103 | 0.722 ± 0.072 | 0.758 ± 0.077 | 0.160 ± 0.050 |
| Non-neuronal | 0.685 ± 0.141 | 0.715 ± 0.139 | 0.783 ± 0.075 | 0.742 ± 0.098 | 0.769 ± 0.113 | 0.470 ± 0.150 |
| Astrocytes | 0.756 ± 0.082 | 0.787 ± 0.097 | 0.794 ± 0.050 | 0.787 ± 0.061 | 0.844 ± 0.063 | 0.070 ± 0.030 |
| Immune Cells | 0.756 ± 0.096 | 0.785 ± 0.097 | 0.782 ± 0.075 | 0.783 ± 0.084 | 0.836 ± 0.099 | 0.040 ± 0.020 |
| Oligodendrocytes | 0.692 ± 0.096 | 0.734 ± 0.118 | 0.771 ± 0.107 | 0.739 ± 0.068 | 0.760 ± 0.112 | 0.310 ± 0.160 |
| OPCs | 0.735 ± 0.064 | 0.751 ± 0.086 | 0.823 ± 0.090 | 0.777 ± 0.046 | 0.810 ± 0.054 | 0.040 ± 0.010 |
| Vasculature | 0.633 ± 0.113 | 0.693 ± 0.111 | 0.666 ± 0.153 | 0.666 ± 0.098 | 0.705 ± 0.106 | 0.010 ± 0.010 |

**Table S3:**
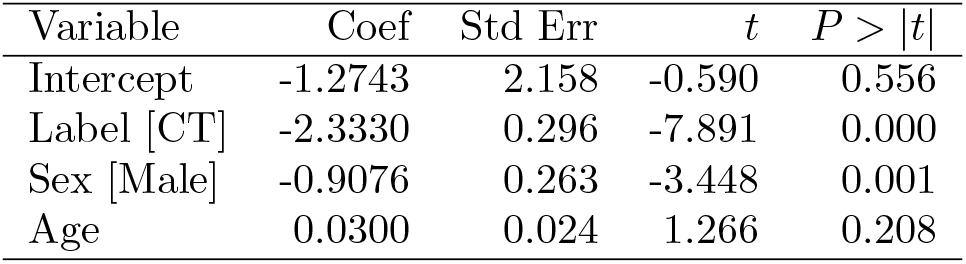
Linear regression performed to assess the association between the first principal component of the sample-level embeddings and age, sex and AD status. Directionality of binary variables is denoted in square brackets.

## References

1. Potter, S. S. Single-Cell RNA Sequencing for the Study of Development, Physiology and Disease. Nature Reviews Nephrology 14, 479–492. issn: 1759-5061, 1759-507X. doi:10.1038/s41581-018-0021-7 (8 Aug. 2018).

2. Travaglini, K. J. et al. A Molecular Cell Atlas of the Human Lung from Single-Cell RNA Sequencing. Nature 587, 619–625. issn: 0028-0836, 1476-4687. doi:10.1038/s41586-020-2922-4 (Nov. 26, 2020).

3. Keren-Shaul, H. et al. A Unique Microglia Type Associated with Restricting Development of Alzheimer’s Disease. Cell 169, 1276–1290.e17. issn: 00928674. doi:10.1016/j.cell.2017.05.018 (June 2017).

4. Neff, R. A. et al. Molecular Subtyping of Alzheimer’s Disease Using RNA Sequencing Data Reveals Novel Mechanisms and Targets. Science Advances 7, eabb5398. issn: 2375-2548. doi:10.1126/sciadv.abb5398 (Jan. 8, 2021).

5. Ravindra, N., Sehanobish, A., Pappalardo, J. L., Hafler, D. A. & Van Dijk, D. Disease State Prediction from Single-Cell Data Using Graph Attention Networks in Proceedings of the ACM Conference on Health, Inference, and Learning ACM CHIL ‘20: ACM Conference on Health, Inference, and Learning (ACM, Toronto Ontario Canada, Apr. 2, 2020), 121– 130. isbn: 978-1-4503-7046-2. doi:10.1145/3368555.3384449.

6. Sehanobish, A., Ravindra, N. & Van Dijk, D. Gaining Insight into SARS-CoV-2 Infection and COVID-19 Severity Using Self-supervised Edge Features and Graph Neural Networks. Proceedings of the AAAI Conference on Artificial Intelligence 35, 4864–4873. issn: 2374-3468, 2159-5399. doi:10.1609/aaai.v35i6.16619 (May 18, 2021).

7. Zeng, F., Kong, X., Yang, F., Chen, T. & Han, J. scPheno: A Deep Generative Model to Integrate scRNA-seq with Disease Phenotypes and Its Application on Prediction of COVID-19 Pneumonia and Severe Assessment http://biorxiv.org/lookup/doi/10.1101/2022.06.20.496916 (2024). Prepublished.

8. Wang, J. et al. scGNN Is a Novel Graph Neural Network Framework for Single-Cell RNA-Seq Analyses. Nature Communications 12, 1882. issn: 2041-1723. doi:10.1038/s41467-021-22197-x (Mar. 25, 2021).

9. Saura, C. A., Deprada, A., Capilla-López, M. D. & Parra-Damas, A. Revealing Cell Vulnerability in Alzheimer’s Disease by Single-Cell Transcriptomics. Seminars in Cell & Developmental Biology 139, 73– 83. issn: 1096-3634. doi:10.1016/j.semcdb.2022.05.007 (Apr. 2023).

10. Mathys, H. et al. Single-Cell Transcriptomic Analysis of Alzheimer’s Disease. Nature 570, 332–337. issn: 0028-0836, 1476-4687. doi:10.1038/s41586-019-1195-2 (June 20, 2019).

11. Kiani, L. Single-Cell Atlas of Alzheimer Disease Vulnerability. Nature Reviews Neurology 20, 505–505. issn: 1759-4758, 1759-4766. doi:10.1038/s41582-024-01008-z (Sept. 2024).

12. Mao, Y. et al. Phenotype Prediction from Single-Cell RNA-seq Data Using Attention-Based Neural Networks. Bioinformatics 40 (ed Kendziorski, C.) btae067. issn: 1367-4803, 1367-4811. doi:10.1093/bioinformatics/btae067 (Feb. 1, 2024).

13. He, B. et al. CloudPred: Predicting Patient Phenotypes From Single-cell RNA-seq in Biocomputing 2022 Pacific Symposium on Biocomputing 2022 (WORLD SCIENTIFIC, Kohala Coast, Hawaii, USA, Dec. 2021), 337–348. isbn: 978-981-12-5046-0 978-981-12-5047-7. doi:10.1142/9789811250477_0031.

14. De Donno, C. et al. Population-Level Integration of Single-Cell Datasets Enables Multi-Scale Analysis across Samples. Nature Methods 20, 1683–1692. issn: 1548-7105. doi:10.1038/s41592-023-02035-2 (11 Nov. 2023).

15. Engelmann, J. P., Palma, A., Tomczak, J. M., Theis, F. J. & Casale, F. P. Mixed Models with Multiple Instance Learning version 2. https://arxiv.org/abs/2311.02455 (2024). Pre-published.

16. Bennett, D. A. et al. Religious Orders Study and Rush Memory and Aging Project. Journal of Alzheimer’s disease : JAD 64, S161–S189. issn: 1387-2877. doi:10.3233/JAD-179939 (Suppl 1 2018).

17. Mathys, H. et al. Single-Cell Atlas Reveals Correlates of High Cognitive Function, Dementia, and Resilience to Alzheimer’s Disease Pathology. Cell 186, 4365–4385.e27. issn: 00928674. doi:10.1016/j.cell.2023.08.039 (Sept. 2023).

18. Wolf, F. A., Angerer, P. & Theis, F. J. SCANPY: Large-Scale Single-Cell Gene Expression Data Analysis. Genome Biology 19, 15. issn: 1474-760X. doi:10.1186/s13059-017-1382-0 (1 Dec. 2018).

19. Stuart, T. et al. Comprehensive Integration of Single-Cell Data. Cell 177, 1888–1902.e21. issn: 00928674. doi:10.1016/j.cell.2019.05.031 (June 2019).

20. Wang, Q. et al. Deep Learning-Based Brain Transcriptomic Signatures Associated with the Neuropathological and Clinical Severity of Alzheimer’s Disease. Brain Communications 4, fcab293. issn: 2632-1297. doi:10.1093/braincomms/fcab293 (Jan. 4, 2022).

21. Gabitto, M. I. et al. Integrated Multimodal Cell Atlas of Alzheimer’s Disease https://www.biorxiv.org/content/10.1101/2023.05.08.539485v1 (2023). Pre-published.

22. Ahern, D. J. et al. A Blood Atlas of COVID-19 Defines Hallmarks of Disease Severity and Specificity. Cell 185, 916–938.e58. issn: 00928674. doi:10.1016/j.cell.2022.01.012 (Mar. 2022).

23. Devlin, J., Chang, M.-W., Lee, K. & Toutanova, K. BERT: Pre-training of Deep Bidirectional Transformers for Language Understanding version 2. https://arxiv.org/abs/1810.04805 (2024). Prepublished.

24. Veličković, P. et al. Graph Attention Networks. Version 3. doi:10.48550/ARXIV.1710.10903 (2017).

25. Brody, S., Alon, U. & Yahav, E. How Attentive Are Graph Attention Networks? Version 3. doi:10.48550/ARXIV.2105.14491 (2021).

26. Kingma, D. P. & Ba, J. Adam: A Method for Stochastic Optimization. Version 9. doi:10.48550/ARXIV.1412.6980 (2014).

27. Hamilton, W. L., Ying, R. & Leskovec, J. Inductive Representation Learning on Large Graphs. Version 4. doi:10.48550/ARXIV.1706.02216 (2017).

28. Wightman, D. P. et al. A Genome-Wide Association Study with 1,126,563 Individuals Identifies New Risk Loci for Alzheimer’s Disease. Nature Genetics 53, 1276–1282. issn: 1061-4036, 1546-1718. doi:10.1038/s41588-021-00921-z (Sept. 2021).

29. Vaswani, A. et al. Attention Is All You Need version 7. https://arxiv.org/abs/1706.03762 (2024). Pre-published.

30. McInnes, L., Healy, J. & Melville, J. UMAP: Uniform Manifold Approximation and Projection for Dimension Reduction. Version 3. doi:10.48550/ARXIV.1802.03426 (2018).

31. Blumenfeld, J., Yip, O., Kim, M. J. & Huang, Y. Cell Type-Specific Roles of APOE4 in Alzheimer Disease. Nature Reviews Neuroscience 25, 91–110. issn: 1471-003X, 1471-0048. doi:10.1038/s41583-023-00776-9 (Feb. 2024).

32. Mrdjen, D. et al. The Basis of Cellular and Regional Vulnerability in Alzheimer’s Disease. Acta Neuropathologica 138, 729–749. issn: 0001-6322, 1432-0533. doi:10.1007/s00401-019-02054–4 (Nov. 2019).

33. Meadows, S. M. et al. Hippocampal Astrocytes Induce Sex-Dimorphic Effects on Memory. Cell Reports 43, 114278. issn: 22111247. doi:10.1016/j.celrep.2024.114278 (June 2024).

34. Lempriére, S. Markers of Vulnerable Neurons Identified in Alzheimer Disease. Nature Reviews Neurology 17, 132–132. issn: 1759-4758, 1759-4766. doi:10.1038/s41582-021-00462-3 (Mar. 2021).

35. Kumari, S., Dhapola, R. & Reddy, D. H. Apoptosis in Alzheimer’s Disease: Insight into the Signaling Pathways and Therapeutic Avenues. Apoptosis 28, 943–957. issn: 1360-8185, 1573-675X. doi:10.1007/s10495-023-01848-y (Aug. 2023).

36. Sharma, V. K., Singh, T. G., Singh, S., Garg, N. & Dhiman, S. Apoptotic Pathways and Alzheimer’s Disease: Probing Therapeutic Potential. Neurochemical Research 46, 3103–3122. issn: 0364-3190, 1573-6903. doi:10.1007/s11064-021-03418-7 (Dec. 2021).

37. Ogunmokun, G. et al. The Potential Role of Cytokines and Growth Factors in the Pathogenesis of Alzheimer’s Disease. Cells 10, 2790. issn: 2073-4409. doi:10.3390/cells10102790 (Oct. 18, 2021).

38. Thakur, S., Dhapola, R., Sarma, P., Medhi, B. & Reddy, D. H. Neuroinflammation in Alzheimer’s Disease: Current Progress in Molecular Signaling and Therapeutics. Inflammation 46, 1–17. issn: 0360-3997, 1573-2576. doi:10.1007/s10753-022-01721-1 (Feb. 2023).

39. Das, N., Raymick, J. & Sarkar, S. Role of Metals in Alzheimer’s Disease. Metabolic Brain Disease 36, 1627–1639. issn: 0885-7490, 1573-7365. doi:10.1007/s11011-021-00765-w (Oct. 2021).

40. Lei, P., Ayton, S. & Bush, A. I. The Essential Elements of Alzheimer’s Disease. Journal of Biological Chemistry 296, 100105. issn: 00219258. doi:10.1074/jbc.REV120.008207 (Jan. 2021).

41. Ejaz, H. W., Wang, W. & Lang, M. Copper Toxicity Links to Pathogenesis of Alzheimer’s Disease and Therapeutics Approaches. International Journal of Molecular Sciences 21, 7660. issn: 1422-0067. doi:10.3390/ijms21207660 (Oct. 16, 2020).

42. Xie, Z., Wu, H. & Zhao, J. Multifunctional Roles of Zinc in Alzheimer’s Disease. NeuroToxicology 80, 112–123. issn: 0161813X. doi:10.1016/j.neuro.2020.07.003 (Sept. 2020).

43. Moynier, F. et al. Copper and Zinc Isotopic Excursions in the Human Brain Affected by Alzheimer’s Disease. Alzheimer’s & Dementia: Diagnosis, Assessment & Disease Monitoring 12. issn: 2352-8729, 2352-8729. doi:10.1002/dad2.12112 (Jan. 2020).

44. Liu, J. et al. APP/PS1 Gene-Environmental Cadmium Interaction Aggravates the Progression of Alzheimer’s Disease in Mice via the Blood-Brain Barrier, Amyloid-β, and Inflammation. Journal of Alzheimer’s Disease 94, 115–136. issn: 13872877, 18758908. doi:10.3233/JAD-221205 (June 27, 2023).

45. Bakulski, K. M. et al. Heavy Metals Exposure and Alzheimer’s Disease and Related Dementias. Journal of Alzheimer’s Disease 76, 1215–1242. issn: 13872877, 18758908. doi:10.3233/JAD-200282 (Aug. 18, 2020).

46. Bakulski, K. M., Hu, H. & Park, S. K. in Genetics, Neurology, Behavior, and Diet in Dementia 813–830 (Elsevier, 2020). isbn: 978-0-12-815868-5. doi:10.1016/B978-0-12-815868-5.00051-7.

